# Co-evolution of ion channels and neurotoxins in cnidarians leads to diversification of ion channel genes

**DOI:** 10.1101/2023.03.11.532225

**Authors:** Anuj Guruacharya

**Author notes:** Correspondence and requests for materials should be addressed to A.G.

## Abstract

Understanding the diversity of ion channels in cnidarians may shed light on the origin and evolution of early nervous systems. It is hypothesized that cnidarian neurotoxins led to the evolution of diverse ion channel proteins in cnidarians. I tested this hypothesis by investigating several evolutionary factors of both cnidarian neurotoxins and their target ion channels. I examined homologs of 250 cnidarian toxins, 75 ion channel genes, and 70 housekeeping genes from 37 transcriptomes/genomes of cnidarian species. Analysis on the presence or absence of genes by species, selection analysis, and evolutionary rate analysis were performed on the homologs of neurotoxin and ion channel proteins. I found evidence of positive selection, correlation between the number of homologous gene families, and difference in the evolutionary rates among the gene families. I have shown for the first time that neurotoxins may have coevolved with the ion channels in cnidarians. This is consistent with an evolutionary arms race between ion channels and neurotoxins leading to extensive diversity of ion channel genes found in cnidarians.

## INTRODUCTION

Cnidarians may provide important clues to the evolution of nervous systems because of their position as one of the earliest diverging lineages of animals exhibiting a rudimentary nervous system (3). Nervous systems allow animals to integrate sensory information and translate this information into behavior. It has been suggested that neurons could have provided early animals with the ability to control a hydrostatic skeleton such as elongating or contracting the body or the ability to open or close feeding appendages according to sensory cues (4). Neurons are also likely to have played an integral role in the evolution of muscle tissue. Yet the origin and evolution of early nervous systems remain obscure (5).

Taxa in the phylum Cnidaria (e.g., sea anemones, hydras, and jellyfishes) have a simple diffuse nervous system unlike the centralized nervous system of virtually all other animals (the Bilateria). Comparative anatomical studies spanning more than 150 years point to the common origin of the nervous systems in the ancestor of Bilateria and Cnidaria, with centralization evolving in the bilaterian lineage. Despite the relative simplicity of their nervous systems, cnidarians have undergone a lineage specific expansion of genes for voltage gated ion channels as shown in Figure 1 (2, 6, 7). Voltage gated ion channels are the primary regulators of ion movement across the membranes of neurons and other excitable cells and are therefore fundamental to action potential formation and signal specificity. Because the expansion of sodium ion channel subtypes in vertebrates appears to correlate with increased neuronal complexity (1), it has been suggested that the expansion of ion channel types in Cnidaria might also correlate with increased neuronal complexity (8).

**Figure 1.**
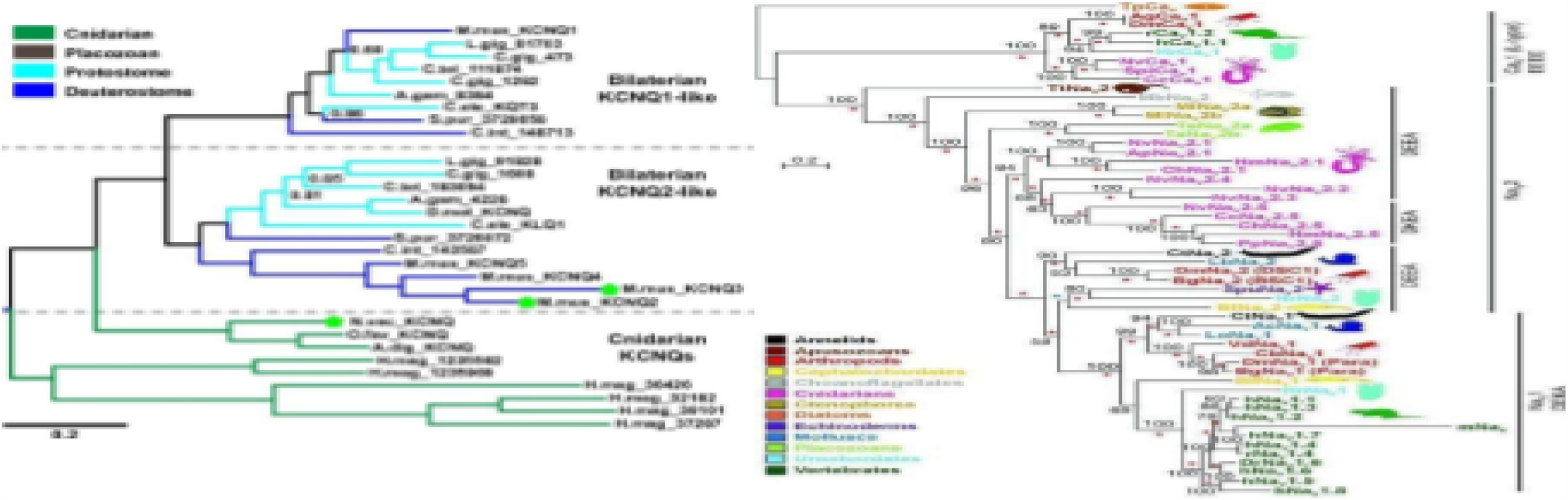
Gene family phylogeny of (left)- KCNQ (K,7) voltage gated potassium channels and (right) sodium ion channels in metazoans. Cnidarians are represented by green color in phylogeny on left and by pink color in phytogeny on top. These figures were reproduced with permission from two groups ADDIN EN.CITE <EndNote><Cite><Author>Barzilai</Author><year>2012</Year><RceNum>3</RecNum< DisplayText>-(1,2) < DisplayText> <record><rec-number>3</rec-number><foreing-keys><key app “EN” Timestamp=“1643912745”>3</key><foreing-keys><ref-type name=“Journul Article”>17<ref-type><contributers><authors><author> Barzilai. Maye Gur <authoer><author>Reitzel. Adam M <authoer><author>Kraus. Jnhanna EM <authoer></author>-Gordon. Dalia</author><author>Tachnau, UIricti </authoer><author> Gurevitz. Michael</authoer><author><contributers><titles><title> Convergent evolution of sodium ion selectivity in metazoan neuronal signaling’<title>Cell reports</secondary-title></titles><periodical><full-title>cell

But given the simple nervous systems of cnidarians, the nature of such possible neuronal complexity is unclear. The evolutionary factors that have driven ion channel diversification in cnidarians, thus, remain poorly understood.

The diversity of ion channels may be related to the broad diversity of toxins that cnidarians produce. Cnidarians are the only phyla possessing nematocysts. These are stinging cells specialized for toxin injection and are considered as part of the nervous tissue (9, 10). Nematocysts inject a cocktail of various peptide and nonpeptide toxins, with neurotoxins being major components of the mix (11). Cnidarian peptide neurotoxins specifically bind to voltage gated ion channels (12), thereby inhibiting nervous system function (13). Sodium channel neurotoxins and potassium channel neurotoxins are the two best characterized toxin groups in these animals (14, 15).

Previous studies have explored phylogenetic analyses of either cnidarian nervous system subunits or cnidarian neurotoxins (16-19), but a systematic investigation of the evolution of both channels and toxins has not yet been reported. With the rise of genomics and the increasing number of cnidarian nucleotide sequences available (20), it has been possible to address questions about ion channel and neurotoxin diversity. Only recently have sufficient cnidarian genome and transcriptome sequences become available to test different hypotheses by examining the homologs of toxins and neural proteins. It has been suggested that evolutionary arms races (21) between predators and prey lead to increasingly potent peptide toxins as well as repeated compensatory changes to defenses against those toxins, causing the evolutionary diversification of both members of interacting protein pairs (22). Well known examples include predator-prey pairs such as grasshopper mice and scorpions (23), and garter snakes and newts (24). Like the coevolving proteins in these predator-prey species, I predicted that ion channels in ancestral cnidarians could have diversified because of natural selection favoring resistance to specific neurotoxins. Diversification of these ion channels may also have been induced by competitive encounters with other cnidarians having a different neurotoxin cocktail than their own.

I thus hypothesize that selective pressure to resist the deleterious effects of neurotoxins may have led to diversification of ion channels in early cnidarians. Such diversification could have involved both nucleotide substitutions as well as gene duplication to expand gene families. If gene duplications have occurred in both toxins and ion channels due to their interactions, then I might predict a correlation between the number of ion channels and neurotoxins genes in different cnidarian species. In addition, if changes in one member of the interacting pair result in compensatory changes in the other, I would expect their evolutionary rates to be correlated along the various lineages. To explore this hypothesis, I acquired genome or transcriptome sequences from 39 diverse cnidarian species. These data were used to construct a species phylogeny of cnidarians which was used to perform gene tree analysis, character analysis on homologous genes, tests of positive selection, and analyses of evolutionary rates of ion channels and neurotoxins.

## MATERIALS AND METHODS

### Data collection

Genome and transcriptome sequences were acquired from 39 different cnidarian species from NCBI genome database, NCBI TSA nucleotide database, and from unpublished transcriptome assemblies from people.oregonstate.edu/∼meyere/data.html (Supplementary Table 1). Genomes of a sponge *(Amphimedon queenslandica)* and fruitfly *(Drosophila melanogaster)* was also obtained for use as outgroups. A set of 70 vertebrate housekeeping genes (25) (Supplementary Table 2) were entered into Uniprot and clustered using a percent similarity identity of 50% to other species via the Uniref50 tool. Only the representative sequence from each cluster was used as the query sequence for blast searches of genomes and transcriptomes. Amino acid sequences for cnidarian venom proteins were collected from venomzone.expasy.org (Supplementary Table 3). Ion channel genes were collected from Uniprot with the GO terms: “sodium ion channel”,” potassium ion channel”,” calcium ion channel”. These channels were then clustered with 50% similarity and only their representative sequences used for further analysis (Supplementary Table 4). The gene families used were of voltage gated sodium, potassium, and calcium channels; actinoporin toxins; Small Cysteine Rich Protines (SCRIPS) toxins; jellyfish toxins; sodium channel neurotoxin type I and type II; and potassium channel neurotoxin type 1, 2, 3, and 5.

**Table 1.** The number of duplications and losses for different gene families in 39 cnidarian species.

| **Gene family** | **Duplication** | **Loss** |
| --- | --- | --- |
| Sodium channels | 5 | 22 |
| Potassium channels | 22 | 56 |
| Calcium channels | 16 | 39 |
| Potassium toxins | 25 | 46 |
| SCRIP toxins | 8 | 15 |
| Actinotoxins | 2 | 21 |

**Table 2.** Results of the selection analyses. LRTs between the respective models for sodium, calcium, potassium, and toxin gene families. The random sites model from CodeML was used to calculate the log likelihoods of individual models. M0, M1, M2, M3, M7, and M8 respectively have a higher number of categories of dN/dS values that are tested out.

| Gene family | Gene name | M0 vs M3 | M1 vs M2 | M7 vs M8 |
| --- | --- | --- | --- | --- |
| Sodium channel | SCN1A | -46.4 | 0 | -125.2 |
| Sodium channel | NALCN | 37.3 | 0 | 0 |
| Calcium channel | CAC1H | 0 | -75.1 | 0 |
| Calcium channel | CAC1B | 131.4 | 0 | -116.1 |
| Calcium channel | CACB2 | 146.8 | 0 | 0 |
| Calcium channel | CAC1C | 355.1 | 0 | 3.9 |
| Calcium channel | CA2D4 | 219.9 | 0.1 | 1.3 |
| Calcium channel | TPC2 | 184.3 | 0 | 3.6 |
| Calcium channel | CAC1I | -5491.9 | 0 | 0 |
| Calcium channel | TPC1 | 0 | 0 | 1 |
| Potassium channel | KCNC1 | 580.6 | 0 | 3.3 |
| Potassium channel | KCNA2 | 391.4 | 0 | 14.6 |
| Potassium channel | KCNA1 | 351.3 | 0 | 14.4 |
| Potassium channel | KCAB2 | 91.6 | 0 | 0 |
| Potassium channel | KCAB1 | 267 | 0 | 1 |
| Potassium channel | KCNH5 | 581 | 0 | 53.9 |
| Potassium channel | KCNH6 | 0 | -5850.2 | 0 |
| Potassium channel | KCNQ5 | 139.3 | 0 | 0 |
| Potassium channel | KCND3 | 229.8 | 0 | 11.3 |
| Potassium channel | CSEN | 71.5 | 0 | 0 |
| Cnidaria small cysteine-rich protein | SCR1 | 121.6 | 0 | 0.9 |
| Dermatopontin | MCTX1 | 72.3 | 1.9 | 8.8 |
| Phospholipase A2 | PA2 | 67.1 | 81.8 | 110.3 |
| Venom Kunitz-type, TypeII Potassium channel toxin | VKT1 | 43.5 | 40.4 | 29.1 |
| Venom Kunitz-type, TypeII Potassium channel toxin | VKT3A | 39.8 | 0 | 0 |
| Venom Kunitz-type, TypeII Potassium channel toxin | VKT3B | 92.9 | 0 | 0 |
| Venom Kunitz-type, TypeII Potassium channel toxin | VKT3C | 46.3 | 0 | 7.9 |
| Venom Kunitz-type, TypeII Potassium channel toxin | VKT3 | 9.2 | 5.6 | 7.1 |
| Venom Kunitz-type, TypeII Potassium channel toxin | VKT3 | 74.9 | 0 | 0.4 |
| Venom Kunitz-type, TypeII Potassium channel toxin | VKT6 | 168.2 | 0 | 0 |

### Species tree

A phylogenetic tree (Supplementary Figure 1) was constructed for all species using the amino acid sequences of 70 housekeeping gene sequences. The 70 housekeeping genes were reciprocally blasted against the genome and transcriptome collection using tblastn and blastx to obtain homologous sequences of each species for each housekeeping gene. The E value used as cut-off for both the blast searches was -10. These genes were then aligned using ClustalO (26), then manually trimmed and aligned again. The genes were then concatenated. PartitionFinder (27) was used to partition the concatenated data into one partition per gene and to find the appropriate evolutionary model for each partition. The LGX model (28) was found to be the best model using Aikai information criterion for all the housekeeping genes. The sponge and fruitfly were included as outgroups. Maximum likelihood was used to find the best phylogenetic tree with RAxML (29). Rapid bootstrapping of 100 replicates was performed using the -a option in RAxML. Mr. Bayes (30) was used to construct the best supported species tree using a Bayesian phylogenetic approach. The rate variation parameter was gamma with 4 rate categories. The chain length for MCMC was 1,100,000 with a subsampling frequency of 200 and a burn in length of 100,000. Treegraph (31) was used to visualize and annotate the species tree with branch lengths and support values.

### Gene tree-species tree reconciliation

In order to assess gene family history, individual gene trees were compared with the species tree. The protein data set was blasted against the genomes/transcriptomes with an E value of -10 using tblastn. They were reciprocal blasted using blastx against the hits obtained. The blast hits for the proteins were grouped according to the gene family they belonged to. The amino acid sequences of the resulting hits were aligned using ClustalW for each protein resulting in 252 multiple sequence alignments. For each gene family, gene alignments were grouped into single file. The best protein evolution model was found using jModelTest (32) for each of the alignments to be Gamma WAG (33). Maximum likelihood gene trees were estimated using RAXML with 100 bootstrap replicates on the potassium channel, sodium channel, calcium channel, potassium toxin and scrip toxin families. Reconciliation analysis used maximum parsimony to estimate the minimum number of gene duplications and losses using Notung (34). An edge weight threshold of 1.4255 was used. Costs/weights were set to 1.5 for duplication, 0 for co-divergence, and 1.0 for losses.

### Gene presence/absence analysis

Correlation analysis was performed based on homologous gene presence or absence data among different species. A character matrix was created using the proteins that were obtained from reciprocal blast for each species. Correlations in the character state matrix were investigated using Pearson’s correlation method in R (35). A linear regression was performed between the total number of channels and the total number of neurotoxins in each species.

### Selection analysis

Gene homologs of the ion channels and neurotoxins were input into Codeml from the PAML package (36) for selection analysis based on non-synonymous substitution rate by synonymous substitution rate (dN/dS) ratios. Several of the genes were present in so few species that selection analyses were not possible. For genes that were present in many but not all species, species trees that were pruned to match the species present were used as the input topologies. A custom script was used to prune the species tree to fit the number of animals in each gene alignment. The alignments were manually curated to verify that they were in frame. I ran random sites models M0, M1, M2, M3, M7, and M8 found in PAML. Selection was inferred from likelihood ratio tests comparing M1 vs. M2 and M7 vs. M8. Likelihood Ratio Tests (LRTs) were performed on M0 vs M3, M1 vs M2, and M7 vs M8. Using a chi square table for one degree of freedom, a cutoff of 3.841 was used to predict statistical significance of positive selection.

### Analysis based on evolutionary rates of gene trees

The dN/dS of the genes that were present in more than four animals were estimated and used to examine evolutionary rates. The mean of the dN/dS of genes in the potassium channel, sodium channel, and potassium toxin families were examined. T-tests were performed to determine if there are significant differences in evolutionary rates between groups.

### Data access

A custom script was built to automate most of the processes mentioned. The custom scripts and related data are deposited in GitHub.

(www.github.com/anuj2054/perseus).

### Methods approval statement

All experiments were performed in sea anemones which are referred to as lower invertebrates.

Therefore, the institutional and/or licensing committee approval was not required.

## RESULTS

### Species tree

Not all of the species I examined have been included in recent phylogenetic analyses (37), thus I conducted a phylogenetic analysis of the currently included species. I used both the Bayesian and maximum likelihood approaches to phylogenetic reconstruction. The topologies produced by both methods were identical (Figure 2), and there were only minor differences in the support values (posterior probabilities and bootstrap values, respectively). Each of the morphologically distinct groups (e.g., anthozoans, myxozoans, hydrozoans) formed highly supported monophyletic groups in the species tree. The myxozoan lineage exhibited substantially longer branch lengths than the rest of the cnidarians which indicates that there is a much higher rate of evolution in that clade.

**Figure 2.**
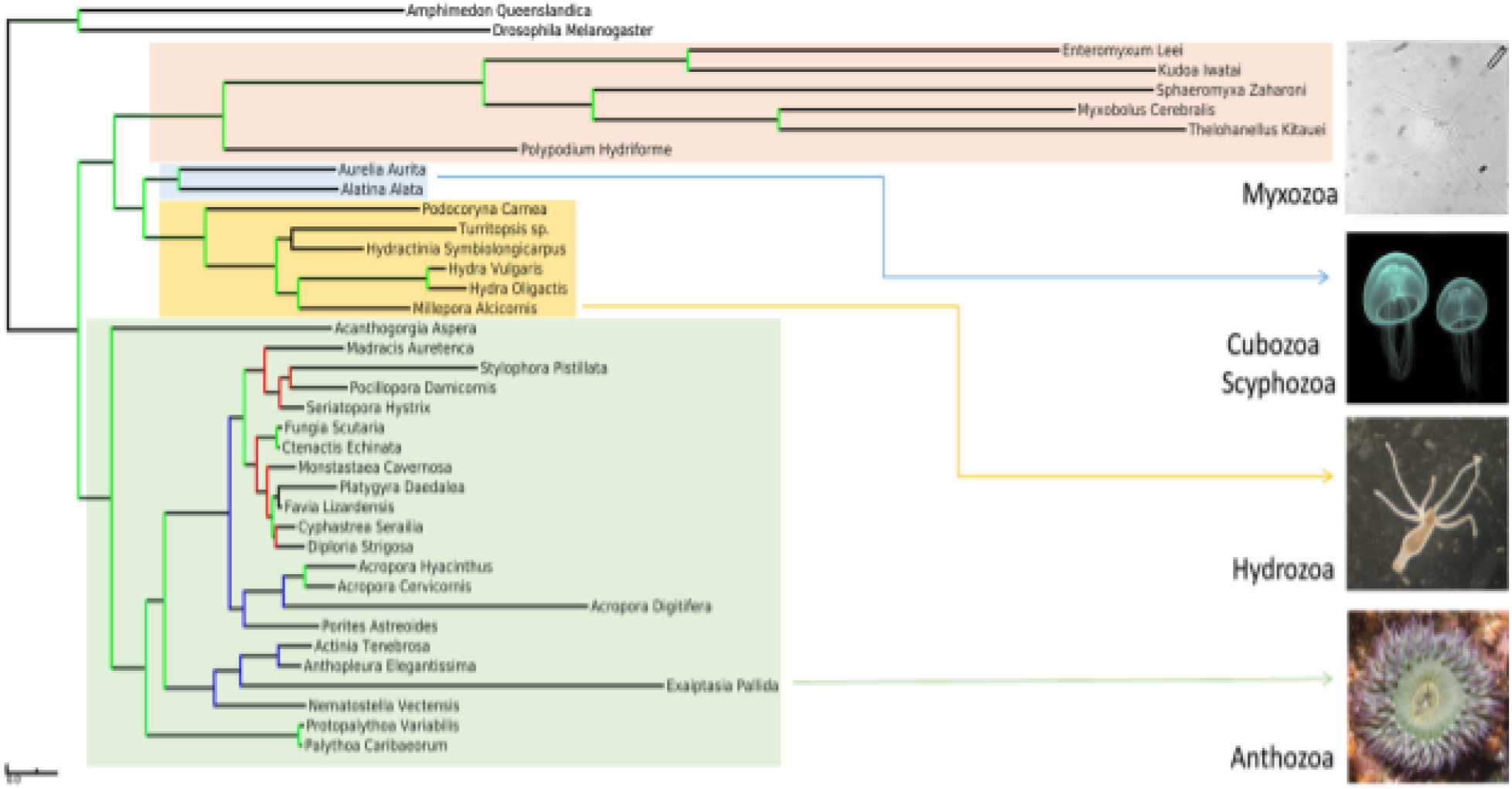
Species tree with branch lengths proportional to substitution rates. The color of node lines Indicates the level of support. Nodes with green lines represent a posterior probability value higher than 0.95 and a bootstrap value higher than 95. Nodes with blue lines represent cases where the posterior probability and bootstrap values were at least 0.90 and 90 and at least one value was less than 0.95 or 95. The nodes with red lines represent nodes where at least one value was less than 0.90 or 90. Each taxonomic class is Indicated by a different background color and Illustrated by a photo of a representative species.

### Occurrence of toxin and ion channel genes among species

The presence or absence of each gene is shown in Supplementary Table 5. Most toxins are found in a restricted number of species. Although the toxin VKTC occurs in 30 of 39 species examined, most other toxins are found in only a few species. Among ion channels, potassium ion channels are found widely in cnidarians, with the major diversity arising from sea anemones. Because of the limited presence of each gene across the whole species tree, many evolutionary comparisons or analyses were limited.

### Gene tree-species tree reconciliation

The gene trees constructed for each of the reciprocal blast hits of the gene families were grouped together into separate alignments. These alignments were used to reconstruct the history of gene duplications and losses. Horizontal gene transfer was assumed not to occur. The results for each of the gene families are shown in Table 1. There was a higher number of duplications and losses observed in the potassium channel family compared to the other ion channel families. Similarly, there was a higher number of duplications and losses in the potassium toxin family compared to the other toxin families.

### Selection analysis

Random site selection analysis was performed on the genes obtained from reciprocal blast. LRTs were performed on the log likelihoods of each gene on the tree under different models of evolution (Table 2). The LRTs were performed on M0 vs M3, M1 vs M2, and M7 vs M8. A positive result for the comparison of M0 vs M3, indicates significant variation in the dN/dS ratio among sites. I would not expect to detect positive selection in any case where there is no significant variation in the dN/dS ratio among sites. A positive result for M1 vs M2 and/or M7 vs M8 provides evidence of positive selection. Ten out of 30 (33%) of toxin or ion channel genes on which the test could be performed exhibited evidence for positive selection. This includes significant evidence of positive selection on four potassium ion channels and three potassium channel toxins. This is the pattern I expect to see if there are evolutionary interactions between toxins and the ion channels that they bind. In addition, one calcium ion channel and 2 other miscellaneous toxins exhibited evidence of positive selection.

### Correlation analysis of gene presence/absence

A table was constructed of the number of homologs of the different types of toxin genes and ion channel genes for each species. The relationship between the number of different ion channel genes and the number of different toxin genes per species was investigated using correlation analyses.

The correlation between neurotoxin and ion channels using the Pearson method was 0.49 with a p value of 0.0046 (not shown). The correlation between potassium toxin and potassium channel using the Pearson method was 0.50 with a p value of 0.0015 (Figure 3). There were too few toxin/channel genes of other types to do correlation analyses by themselves.

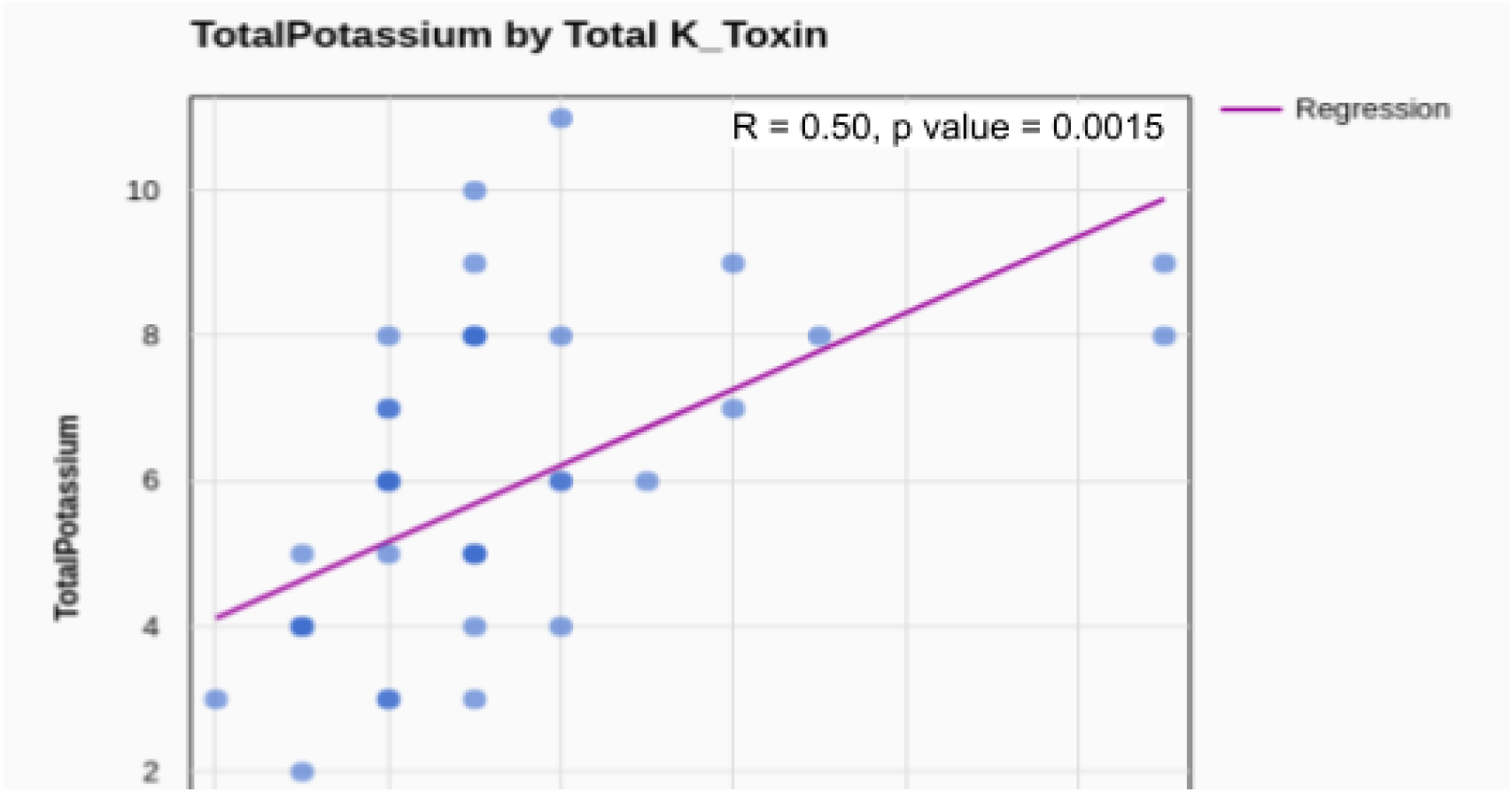

### Analysis of evolutionary rates

The dN/dS of the genes that were present in more than four animals were estimated and used to examine evolutionary rates. The mean of the dN/dS (shown in Figure 4) of the gene related to potassium channels, sodium channels, and potassium toxins were examined as these were the only groups containing specific genes found in at least four taxa. T-tests were performed to determine if there are significant differences in evolutionary rates between groups. The p value of calcium channels vs. potassium channels was 0.3626. The p value of potassium channels vs. potassium toxins was 0.07561. The p value of calcium vs. toxin was 0.0731. Thus, there were no significant rate differences between any class of genes, but the evolutionary rates of toxin genes were appreciably higher than rates of the ion-channel genes.

**Figure 4.**
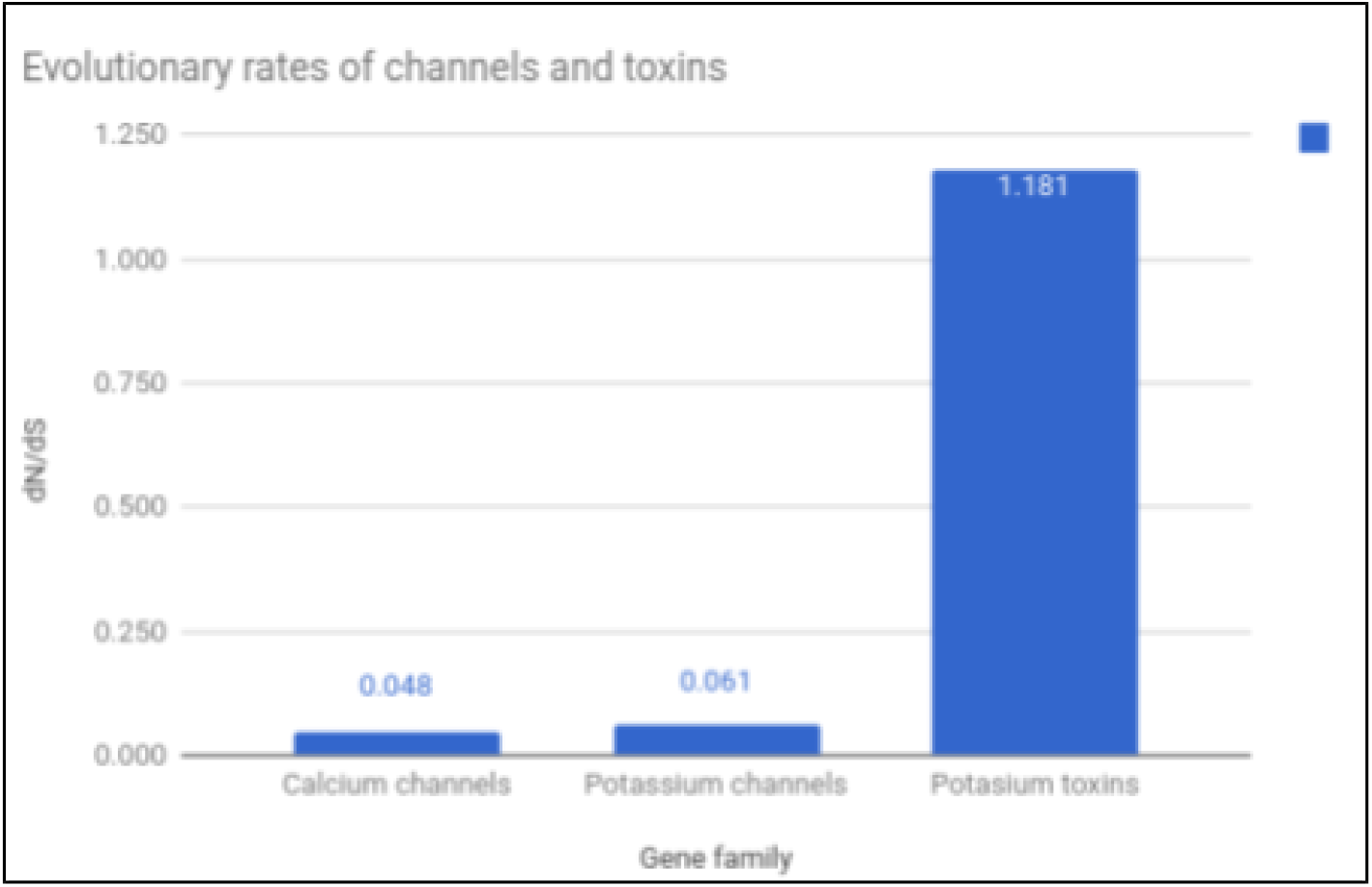
Evolutionary rates of channels and toxins. The mean of the dN/dS evolutionary rates of the calcium channels, potassium channels and potassium toxins. These values were obtained from using Model 6 In CODEML.

## DISCUSSION

My results provide a novel perspective on the evolution of neurotoxins and their target voltage gated ion channels in Cnidaria. The level of ion channel diversity observed in cnidarians, at both the gene family and nucleotide levels, is not observed in any other animal group. The co-occurrence of this diversity with the diverse set of toxins unique to cnidarians suggests a causal relationship. Moreover, the greatest diversity of ion channels at the intraspecific level occurs in sea anemones, with a correspondingly high diversity of toxins, while the lowest diversity of ion channels and neurotoxins occurs in the parasitic cnidarians. This pattern also holds for comparisons among whole clades, e.g., taxonomic classes, (although it is possible this could be a function of the level of taxonomic sampling). These observations reinforce the evolutionary relationship between neurotoxins and their ion channel targets in Cnidaria and because voltage gated ion channels are integral components of neurons, this has important implications for nervous system evolution in early animals. Conclusions of this study require a few caveats. One limitation is in the datasets themselves. In many cases I used transcriptome sequences. A transcriptome only contains sequences of those genes that have been expressed at the time the tissue is sampled. To reduce this problem, only transcriptomes whose source tissue contained tentacular tissue were used in this study. Another limitation involves the sensitivity of reciprocal blast to identify short sequences (as in the case of toxin genes). Shorter sequences have a lower sensitivity of being detected than longer sequences (38). For short sequences, proteins having related functions may not show overall high similarity yet contain a few short amino acid motifs or residues that are highly conserved. Alternatively, for long sequences, proteins may show overall high percentage similarity but can contain a few differences in important functional domains that change their function (39). Using Hidden Markov Model profiles was not an appropriate approach since it requires a protein sequence database of each animal. For proteins with short amino acid sequences, like the neurotoxins, translating them from assembled nucleotide contigs in a transcriptome to proteins sequences leads to a high number of false positives. Finally, I was not able to perform many analyses due to the limited presence of many genes across taxa. For example, the sparse distribution of specific homologs among taxa meant that in many cases there was not a sufficient number of sequences to perform tests of positive natural selection. In addition, I had hoped to perform analyses of evolutionary rate correlations among genes, yet because many homologs were absent from most taxa, there was not sufficient data to perform correlation analysis. In the end, I believe the analyses that I was able to perform provide sufficient support for an evolutionary relationship between neurotoxins and ion channels.

### Rapid evolution of parasitic cnidarians

The Myxozoa present a unique situation so I will discuss them separately. The general evolutionary pattern observed for parasitic cnidarians is one of rapid evolution with a small number of ion channels and neurotoxins. Two common reasons for rapid evolution are shorter generation times or a change in environment (40). It appears that the environment of parasitic cnidarians would have changed substantially with a change in lifestyle. This dramatic change in addition to adaptation to a range of hosts (e.g., from jellyfish to salmon) would have required rapid and extensive changes at a genomic scale.

A low number of neurotoxins and their target ion channels in parasitic cnidarians could be due to a lack of necessity to attack prey, defend against predators, or compete for space. As parasites acquire energy from their hosts, killing or paralyzing prey using neurotoxins is not necessary. Even though these animals retain nematocysts their utility remains unclear. For example, whirling disease is a common disease found in salmonid fishes that are caused by parasitic cnidarian, *Myxobolus cerebralis* (41). In this case the animal lives in the cartilage and bone of the fishes and are known to cause neurological damage (42, 43). The role of nematocysts and their neurotoxins as part of the infection process or during larval stages remains unknown. However, myxozoans have been reported to have undergone an extreme reduction in genome size and gene content (44), thus many neurotoxin and ion channel genes may have been lost since their divergence from free living ancestors.

### Ion channels and neurotoxins may have coevolved in an evolutionary arms race

Evolution of ion channels and their toxins is a dynamic process with the hallmarks of an evolutionary arms race. I have provided several lines of evidence that are consistent with this model. One is that there clearly have been numerous gene duplications along with two to four times as many gene losses. Potassium ion channels and potassium channel toxins are the most diverse systems with the highest number of duplications and losses. If ion channels are evolving in response to the toxins that bind to them, a diversity of channels might arise as a result of an increasing number of toxins. Thus, duplicated and diverged (45) channels could provide an escape mechanism to allow species to evade the toxins of related taxa. Toxin gene expansion has been observed on various branches of the cnidarian tree. The very short (mostly close to 100 amino acid) length of common toxin genes suggests there may be a limited number of effective forms, increasing the chance of convergent toxin evolution. Nonetheless, the loss of toxin genes appears to be far more common, suggesting that many toxin genes may be lost after alternative forms arise. The independent gain and loss of the toxins as indicated by the gene tree species tree reconciliation studies suggests a very dynamic positive feedback system where toxin genes evolve in response to corresponding evasive changes in their target ion channel genes.

The second line of evidence supporting the hypothesis is that I observed a significant correlation between the number of toxin genes and the number of ion channel genes within taxa. This is consistent with a scenario where multiple alternative ion channels arise in response to diversifying toxins. This correlation suggests that neurotoxins may have been an important driving force for the evolution of neural and muscular systems in which ion channels play an important role. While correlation does not indicate causation, this is additional evidence that is consistent with a toxin-channel evolutionary arms race.

The third line of evidence involves the results obtained from analysis of nucleotide sequence (codons) of specific genes. There is evidence for positive natural selection on several ion channel and toxin proteins (among those that were present in a sufficient number of taxa to perform tests of selection). This is particularly true for potassium channels and the toxins that target them. This is exactly the result I would expect if toxins and their target ion channels are co-evolving. There may well be selection on other genes in my study, as the method I used is known to be conservative in cases where dN/dS ratios are elevated but still less than 1.0 (46). I note that recent reports have concluded that venoms from evolutionarily younger lineages such as snakes and cone snails were under positive selection, while more ancient lineages such as cnidarians, spiders, centipedes and scorpions tended to be more constrained under negative selection (16). Yet they suggested that episodic bursts of adaptive selection could occur on most toxin types with shifts in ecological parameters. I suggest that many cnidarian toxins have undergone positive selection due to recent arms race interactions with their associated ion channels.

In many arthropods venom is used as a weapon in predation as well as for intraspecific competitive interactions (47). Cnidarians similarly use venom for predation and defense (48, 49). Many anemones as well as scleractinian corals are known to use venom to attack other individuals (Nelsen et al, 2014; Williams, 1991). They attack conspecifics or related taxa in competitive interactions to protect or expand their territories (50, 51). Thus, competition could have led to the diversification of the neurotoxin arsenal and the need for protection against the arsenals of conspecifics and close relatives.

Here I provide the first integrated analyses of cnidarian neurotoxins and their voltage gated ion channel targets. My results provide multiple lines of evidence that consistently support the coevolution of neurotoxins and ion channels in cnidarians. The evolutionary arms race scenario I have described provides a compelling explanation for the unique diversity of ion channels and toxins found in cnidarians. This study has important implications for the study of evolution of early nervous systems which is of increasing importance in evolutionary biology. In addition, the coevolution of toxins and ion channels provides a foundation for further studies of nervous systems as more taxa and genomes become available.

## Consent for publication

Not applicable

## Supporting information

https://docs.google.com/spreadsheets/d/1jVV5hEDfCCihZBUzbcGjw7nXJL3kd6Mj/edit?usp=sharing&ouid=104221907928556892076&rtpof=true&sd=true

## Availability of data and materials

Data and materials are available in the main manuscript as well as in the supplementary information.

## Competing interests

The Author declares that there is no conflict of interest.

## Funding

This research received no specific grant from any funding agency in the public, commercial, or not-for-profit sectors.

## Authors’ contributions

A.G. performed, designed the study and performed statistical analysis. A.G. wrote and approved the final manuscript.

## Acknowledgements

Author would like to thank the University of Oklahoma for providing a safe space for research.

## Notes

### Competing Interest Statement

The authors have declared no competing interest.

